# From peer-reviewed to peer-reproduced: a role for data standards, models and computational workflows in scholarly publishing

**DOI:** 10.1101/011973

**Authors:** Alejandra N. González-Beltrán, Peter Li, Jun Zhao, Maria Susana Avila-Garcia, Marco Roos, Mark Thompson, Eelke van der Horst, Rajaram Kaliyaperumal, Ruibang Luo, Tin-Lap Lee, Tak-wah Lam, Scott C. Edmunds, Susanna-Assunta Sansone, Philippe Rocca-Serra

**Author notes:** contributed equally to the work (Alejandra N. González-Beltrán, Peter Li, Philippe Rocca-Serra). & (Susanna-Assunta Sansone, Philippe Rocca-Serra).

## Abstract

**Motivation:** Reproducing the results from a scientific paper can be challenging due to the absence of data and the computational tools required for their analysis. In addition, details relating to the procedures used to obtain the published results can be difficult to discern due to the use of natural language when reporting how experiments have been performed. The Investigation/Study/Assay (ISA), Nanopublications (NP) and Research Objects (RO) models are conceptual data modelling frameworks that can structure such information from scientific papers. Computational workflow platforms can also be used to reproduce analyses of data in a principled manner. We assessed the extent by which ISA, NP and RO models, together with the Galaxy workflow system, can capture the experimental processes and reproduce the findings of a previously published paper reporting on the development of SOAPdenovo2, a de novo genome assembler.

**Results:** Executable workflows were developed using Galaxy which reproduced results that were consistent with the published findings. A structured representation of the information in the SOAPdenovo2 paper was produced by combining the use of ISA, NP and RO models. By structuring the information in the published paper using these data and scientific workflow modelling frameworks, it was possible to explicitly declare elements of experimental design, variables and findings. The models served as guides in the curation of scientific information and this led to the identification of inconsistencies in the original published paper, thereby allowing its authors to publish corrections in the form of an errata.

**Availability:** SOAPdenovo2 scripts, data and results are available through the GigaScience Database: http://dx.doi.org/10.5524/100044; the workflows are available from GigaGalaxy: http://galaxy.cbiit.cuhk.edu.hk; and the representations using the ISA, NP and RO models are available through the SOAPdenovo2 case study website http://isa-tools.github.io/soapdenovo2/. **Contact:**

## Introduction

Several reports have highlighted the practical difficulties in reproducing results from published experiments [1–4]. That a basic tenet of scientific research cannot be fulfilled has fuelled growing concerns from stakeholders with an acute interest in scientific reproducibility such as universities, industry, funding agencies, the wider research community as well as the public. A failure to reproduce published scientific findings adversely affects scientific productivity and, in worse cases, may lead to retraction [5]. Moreover, it casts doubt on the quality of the peer-review process. Therefore, publishers have renewed efforts to mitigate the shortcomings of the science reported in their journals. Amongst the incentives tried by publishers are the lift on restrictions on the length of methods sections, the creation of data publication platforms, such as GigaScience^1^ and Scientific Data^2^, the provision of a statistical review of numerical results where appropriate and the requirement for data to be deposited in open-access repositories. These efforts have in part been driven by position statements from funding agencies, publishers and researchers advocating more widespread data sharing^3^.

Computational frameworks and data models now exist which can be used to structure scientific data and their analyses. In this article, we investigate three conceptual community data models for providing structured reporting of findings and scientific workflows for capturing the data analysis pipeline. Investigation/Study/Assay (ISA) is a widely used general-purpose metadata tracking framework with an associated suite of open-source software, delivering rich descriptions of the experimental condition information [6]. The ‘Investigation’ provides the project context for a ‘Study’ (a research question), which itself contains one or more ‘Assays’ (taking analytical measurements and key data processing and analysis steps). The transformations of data underlying an analysis can be represented as steps within a scientific workflow that can be automatically executed and repeated on platforms such as Taverna [7] and Galaxy [8]. Nanopublication (NP) is a model which enables specific scientific assertions to be annotated with supporting evidence, published and cited [9]. Lastly, the Research Object (RO) model enables the aggregation of the digital resources contributing to findings of computational research, including results, data and software, as citable compound digital objects [10]. Combined, these conceptual models facilitate the validation of the findings and assist the reuse and understanding of the results.

Our study addresses the question of whether such data and workflow representation frameworks can be used to assist in the peer review process, by facilitating evaluation of the accuracy of the information provided by scientific articles with respect to their repeatability. We use the ISA framework, the Galaxy workflow platform, NP and RO models on an article in GigaScience. Jointly published by BioMed Central and BGI, GigaScience is linked to a database, GigaDB [11], hosting large scale datasets, but also scripts used to analyse a dataset associated with the publications. The article [12] was selected on the basis that all the data, the analysis scripts used and extensive documentation were all publicly available in GigaDB^4^. However, as we will show, even deposition of the data and the software required to perform the analysis in an open repository does not guarantee reproducibility. Even though seven referees had tested a number of the data sets and analysis scripts^5^, we found issues with reproducing the actual results published in the article. In this paper, we show how the combination of data and workflow representation models play a crucial part in highlighting important experimental elements, otherwise easily missed, and enhance data reporting, data review and data publication processes.

## Results

### 0.1 SOAPdenovo2 experiment overview

The article by Luo *et al* [12] describes the development of SOAPdenovo2 and its evaluation as a computational tool for the *de novo* assembly of genomes from small DNA segments read by next generation sequencing (NGS). Improvements were made at each step of the de Bruijn graph-based algorithm implemented by SOAPdenovo1. This new algorithm was evaluated against four NGS data sets from two bacterial genomes (*S. aureus* and *R. sphaeroides*), one insect genome (*B. impatiens*) from the Genome Assembly Gold-standard Evaluations (GAGE^6^) competition [13], and the human YH Asian Genome data set [14]. The performance of SOAPdenovo2 was compared with its predecessor, SOAPdenovo1 [15], and ALL-PATHS-LG [16].

### 0.2 Reproducing the results from the paper with Galaxy workflows

Our reproducibility effort focused on developing Galaxy workflows, re-creating the data analysis processes used in calculating the results in Tables 2, 3 and 4 in [12], which show the performance of SOAPdenovo2 in assembling the four genomes aforementioned. Prior to developing the workflows, SOAPdenovo2, its pre- and post-processing tools had to be integrated into a Galaxy server^7^ using their command-line interfaces. These were then combined within Galaxy workflows, thus recapitulating the computational steps the SOAPdenovo2 authors used in *bash* and *perl* scripts for assembling the genomes and evaluate the performance of their new assembler^8^. Due to both insect and human data sets’ large sizes, we were not able to develop executable workflows for assembling these genomes as our public server could not meet the memory needs of up to 155 GB, as indicated by the SOAPdenovo2 authors for building the human genome.

Galaxy workflows were developed to assemble the genomes for *S. aureus* and *R. sphaeroides*. However, for those genomes, two additional steps that were not found in the authors’ *bash* scripts were required to reproduce the statistics. A step was required to break scaffolds between any gaps into separate sequences. Another was needed to calculate the actual genome assembly statistics in Table 2 from [12], performed by an analysis script^9^ developed for use in the GAGE genome assembly competition [13]. Both of these steps were added to the SOAPdenovo2 genome assembly Galaxy workflows for *S. aureus* and *R. sphaeroides*. The results obtained from the execution of these workflows were almost identical to those published in [12] and are available in Table 1. By deploying SOAPdenovo1 and ALL-PATHS-LG [17] as tools within Galaxy, it was possible to re-implement genome assembly and reproduce the results from [12], albeit with minor discrepancies (see Table 1).

**Table 1.** Results from reproducing Table 2 of the original paper, where the original results are shown in between parenthesis.

| Species | Algorithm | Contig |  |  |  | Scaffold |  |  |  |
| --- | --- | --- | --- | --- | --- | --- | --- | --- | --- |
|  |  | Number | N50(kb) | Errors | N50 corrected (kb) | Number | N50(kb) | Errors | N50 corrected (kb) |
| S. aureus | SOAPdenovo1 | 79 | 148.6 | 156 | 23 | 49 | 342 | 0 | 342 |
|  | SOAPdenovo2 | 80 | 98.6 | 25 | 71.5 | 38 | 1086 | 2 | 1078 |
|  | ALLPATHS-LG | 37 | 149.7 | 13 | **119 (117.6)** | **11** | 1477 | 1 | 1093 |
| R. sphaeroides | SOAPdenovo1 | **2241 (2242)** | 3.5 | **400 (392)** | 2.8 | 956 | **106(105)** | **24(18)** | **68 (70)** |
|  | SOAPdenovo2 | 721 | 18 | 106 | 14.1 | 333 | 2549 | 4 | 2540 |
|  | ALLPATHS-LG | 190 | 41.9 | **30(31)** | 36.7 | 32 | 3191 | 0 | **0 (3310)** |

**Table 2.**
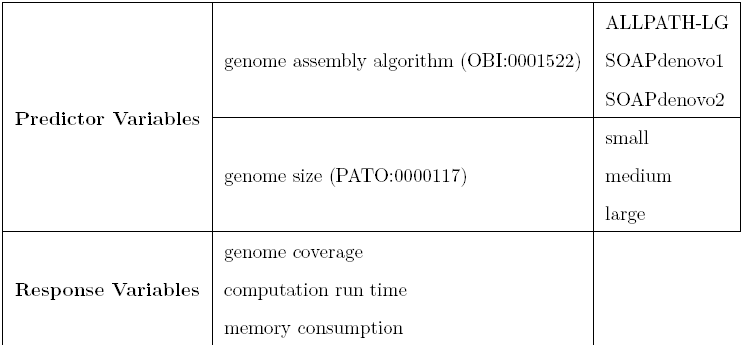
Predictor and response variables for the SOAPdenovo2 study, as identified in the ISA-TAB documents

|  |  |  |
| --- | --- | --- |
| **Predictor Variables** | genome assembly algorithm (OBI:0001522) | ALLPATH-LG<br>SOAPdenovo1<br>SOAPdenovo2 |
|  | genome size (PATO:0000117) | small<br>medium<br>large |
| **Response Variables** | genome coverage<br>computation run time<br>memory consumption |  |

### 0.3 Modeling the experimental process using ISA

The ISA research object provides constructs to describe study design and experimental variables. It can accommodate minimal information guidelines [18], which may insist on reporting such information. As indicated in the experiment overview, in [12], four genomes from three distinct phyla, representing 3 points along a genome size gradient covering several orders of magnitude, were used to test 3 genome assembly software. Thus, we summarized the experiment as a 3×3 factorial design, with two independent variables or factors declared: software and genome size – cast as `Study Factors` in the ISA syntax. For both variables, three discrete levels were found and reported in a `Factor Value` field in an ISA assay table. As [12] compares *de novo assembler* methods, the independent variable levels do not affect the samples and do not need to be reported at the study table level.

Next, we represented the data points, or members of each study group. As [12] accounts for refinements to the first published diploid genome sequence of an Asian individual (referred to here as *Chinese Han genome* or *YH genome*) [14, 19] with new reads generated on the newer Illumina platform, the assay template *”genome sequencing using nucleotide sequencing”* was chosen from the various wet lab workflow templates available from ISAcreator, the curation tool in the ISA infrastructure. This ensures meeting annotation requirements covering key steps of specific experimental processes, enabling direct deposition to the European Nucleotide Archive [20] through ISAcreator or to the Short Read Archive repository [21]. This representation allows distinguishing newly generated data from downloaded data when declaring inputs in the genome assembly processes.

The ISA model minimal implementation guidelines instruct to systematically report data file and software locations as resolvable identifiers. The guideline resulted in detecting missing files (for an example, refer to Table 2) and unresolvable file references. It also revealed a lack of unambiguous identification of the reference genomes used to perform the alignment step. We achieved a resolution through direct communication with the authors of [12], clarifying that the NCBI human reference genome *hg19* mentioned in [12], known to GenBank as Genome Reference Consortium Human Build 37 (GRCh37)^10^, corresponds to GenBank Assembly *ID:GCA 000001405.1*. This fact allowed to disambiguate the reference genome with its subsequent releases (7 in total).

We then focused on identifying the response variables, and their units, used to assess assembly software efficiency. Information from result tables in [12] was extracted, identifying six metrics: i.) genome coverage (as a percentage), ii.) contig N50, iii.) scaffold N50 (stated in kb or base pairs (bp)), iv.) number of errors, v.) run time (stated in hours) and vi.) peak memory usage (stated in gigabytes). For each response variable, Table 1 in supplementary material collates definitions as reported in [16]. The first four metrics provide estimates on assembly efficiency and accuracy, whilst the last two give insights into computational efficiency and therefore depict the savings the most efficient computational method can offer in terms of time and memory. Correspondence with the original authors confirmed that all metrics were calculated using an analysis script from GAGE [13], executed in a fixed environment on each of the genome assembly software output files, thus guaranteeing protocol consistency. Using ISA, sequence analysis and software comparison outputs were reported relying on `Derived Data File` fields used to supply file paths or Gigascience document object identifiers (DOIs) to relevant manuscript objects.

The ISA representation of the study by [12] is released as an ISA-Tab archive and a semantic representation using the Resource Description Framework (RDF) [22]. The latter relies on the *linkedISA* software component [23], using a mapping to Open Biological and Biomedical Ontologies (OBO) resources [24]. In particular, mapping to the Ontology for Biomedical Investigations (OBI) [25] ensure interoperability with several projects using OBI and alignment with ISA configurations. In addition, linkedISA can also be configured with additional mappings and for the conversion of the SOAPdenovo2 experiment, we used a mapping to the provenance ontology (PROV-O^11^).

### 0.4 Publishing findings as Nanopublications

First, we considered following key findings to be expressed as nanopublications:

1. genome coverage increased and memory consumption was 2/3 lower (during the point of largest memory consumption) over the human data when comparing SOAPdenovo2 against SOAPdenovo1.
2. improvements in contig and scaffold N50 metrics when considering SOAPdenovo2 versus SOAPdenovo1 for *S. aureus*, *R. sphaeroides* and YH dataset, as presented in Tables 2 and 4 of [12]

These key findings were extracted from the abstract and main conclusions of the article. We tracked the provenance of the statements by identifying the corresponding rows in the tables of the article [12] and complemented them with more statements extracted from those tables, taking into account the response variables, as identified earlier. This process resulted in 9 assertions that were embedded into 9 nanopublications.

The nanopublications were created following a novel methodology that combine OntoMaton [26] and NanoMaton^12^ software tools. Collected statements were structured as triples in a Google spreadsheet, using the OntoMaton widget, a component of the ISA software suite [26] that accesses community ontologies portals [27, 28]. The collaborative environment allowed review, discussion and incremental improvement until satisfactory expressivity and clarity was reached. The statements were processed with the NanoMaton software component, which converts the OntoMaton templates in RDF. A conceptual overview of how ISA and a nanopublication are related is presented in Figure 1.

**Figure 1.**
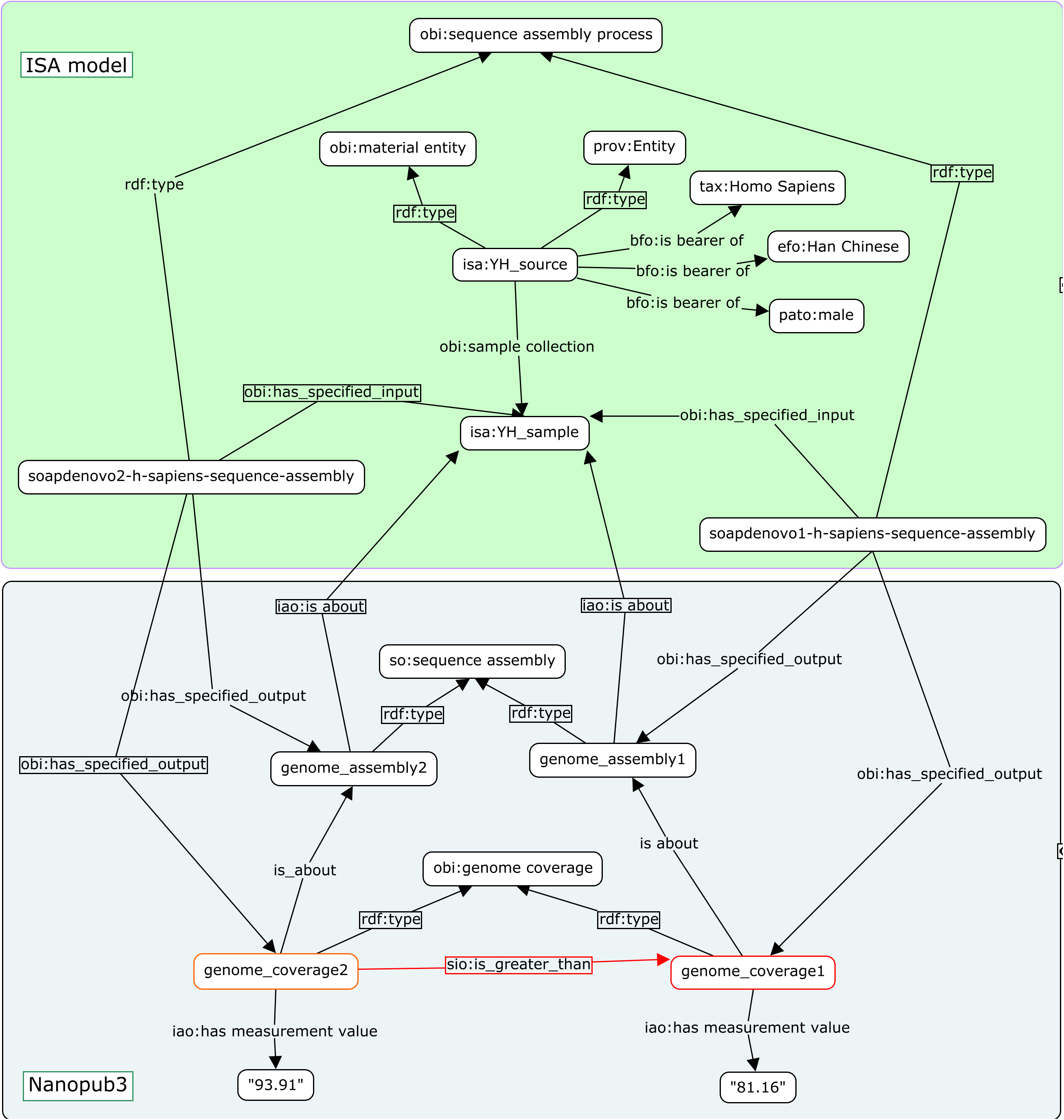
A diagram showing part of the ISA experimental description and a nanopublication relying on elements of the ISA representation.

The nanopublication guidelines advocate the use of existing semantic types to create NPs as Linked Data. We thus relied on OBI [25], STATO^13^ and SIO [29] but none provided semantic types for response variables such as *genome coverage*, *random access memory*, *computation runtime* that were used in [12]. Gaps in domain coverage is a known caveat in the Semantic Web approach, especially when carrying out de-novo semantic modelling. It either requires issuing a term request in existing resources or creating a new ontology. We chose the latter owing to familiarity with OBI procedures, thus ensuring rapid processing and the completion of nanopublications seamlessly consistent with the linkedISA RDF representation.

The provenance component of the NP model also required special attention. Since the NP translates a written statement from a manuscript into RDF triples, it inherently retains an interpretative aspect by those formulating it. We therefore included all contributing parties, from the original authors [12] to the *semantic translators* who crafted the NPs.

### 0.5 Preserving computational workflows and aggregating ISA and NP representation as Research Objects

The ISA framework establishes scientific rigor by requesting scientists to report their experiment results along with information about the data analysis process and experiment design. The Research Object model [10] advocates the sharing of information related to a study as one structured aggregation object, in order to facilitate the validation of the findings and assist the reuse and understanding of the results. The use of this model should be guided by some minimal principles, including: 1) making the aggregation object as well as every one of its component identifiable, so that they can be referred to, shared, and cited; 2) associating basic metadata with an RO and its resources, including how they relate to each other, so that they can be interpreted; 3) associating basic provenance information with the RO and its resources, like where they came from, when *etc.*, to assist attribution, versioning, citation, and reproducibility; and 4) wherever applicable, soliciting rules for assessing the completeness of an RO, i.e. containing all things required for the purpose of creating this RO. In this study, these principles were applied to capture resources related to the Galaxy workflows created by GigaScience for generating the results presented in Table 2 in [12]. The RO thus contains links to the input data used by the workflow, the Galaxy workflow itself (made available through the export function of the GigaScience Galaxy instance) and the provenance statements about the inputs used. Everything in this RO as well as the RO itself is uniquely identified and can be referred to. A list of 5 rules was implemented as a checklist [30] to assess the completeness of the RO.

We observed some redundancy in capturing workflow inputs and outputs by the workflow-centric RO and the linkedISA conversion [23]. One could in fact envisage a reuse of RO workflow extension in an non-BFO based linkedISA conversion of ISA-Tab document.

### 0.6 ISA, NP and RO models: a complementary set of representational resources

We have described how these models have been harnessed to represent a computational experiment comparing genome assembler efficiency [12]. In order to convey the information payload held in each of the components more clearly, the SOAPdenovo2 case study website^14^ includes a table summarising a number of query cases and highlights which model allows those queries to be answered. The overall study is described in an ISA-Tab document then converted to an RDF representation by the linkedISA software. At the other end of the spectrum, key experimental findings have been expressed as nanopublications, a semantic web compatible representation of the most salient results. All of these representations are placed in a broader context through a wrapper layer realised in the form of a Research Object.

## Discussion

The authors of the original SOAPdenovo2 paper [12] strived to make their work reproducible, making their source data, tools and scripts all accessible together with documentation. Yet, it still took about half a man-month worth of resources to reproduce the results reported in Table 2 from [12] in Galaxy workflows. This indicates that it is not a lack of contributions or efforts by authors that hampers reproducible research practice. Rather, it is a lack of understanding of what needs to be provided to make this vision materialize. It also identifies a need to develop instructions for authors that go beyond traditional narrative papers. Our work revealed several distinct reasons leading to reproducibility collapse, even when software and data are available. These can be cast into the following categories:

i. ambiguities in resource identification,
ii. absence of computer readable descriptions of inputs, computational workflows and outputs,
iii. absence or limitation of available computational resources,
iv. absence of identification of main elements in terms of experimental design such as predictor and response variables
v. limitation of depth and breadth of semantic artefacts, disambiguating unclear meaning of experimental elements.

So more generally, it is about dealing chiefly with ambiguity about experimental planning, followed by incompleteness in accounts.

Garijo *et al* [31] outline some desiderata and guidelines to authors to improve the reproducibility of their results. Their paper focuses on reproducing computational results and the desiderata and guidelines emphasise on making available the input data, a data flow diagram, the software and its configurations together with intermediate data. The article we chose for this case study [12] does comply with most of the guidelines: the authors provided *bash* and *shell* scripts together with documentation and indications on how to obtain the input data. While intermediate data was not available, it could be obtained by running the given scripts. According to the classification provided by Garijo *et al* [31], the results could be reproduced by a novice user and by following the documentation, it was possible to reproduce the results with Galaxy workflows with minimum interaction with the authors. The GigaGalaxy platform now provides all the facilities, including workflow definitions and intermediate data, to re-enact the execution and reproduce the results. However, we identified other issues hampering reproducibility, which we describe later in this section.

But when considering the calls for reproducibility, let’s analyse its costs and who should bear them. *De novo* assembly of large genomes requires significant computational resources. Allowing for re-enacting those processes has an obvious economical footprint which very rapidly places a cap on what can currently reasonably be offered. Typically, using an Amazon Web Services (AWS) instance ‘cr1.8xlarge’ with up to 244GB memory suited for repeating the large genome assembly, costs USD 3.5 per hour^15^. Repeating the YH genome assembly thus represents a USD 200-300 expenditure, excluding unavoidable storage cost. This raises the question *who most critically needs to reproduce all publication results?* Presumably, reviewers and journal editors should be the primary beneficiary of this attention. It is evident that not all results can be re-enacted owing to the associated operational costs, however, it is a pragmatic position to require that viable alternatives be provided to enable evaluation and review in order to establish trust in the results. This is the approach chosen by GigaScience.

The attention therefore shifts to certain qualitative aspects associated to the reporting of scientific experimentation. Despite the *big data* hype and associated controversial claims [32,33], for most scientists, either computational or bench biologists, dispensing with the theory or with experimental designs is not an option. We show how to make the most of this information to perform a deeper review and help produce better reports.

The simplest issue to address when improving experimental reporting is resource identification. It constitutes our first and easiest recommendation: **unambiguously identify electronic resources, such as records downloaded from public repositories, by providing their official identifiers**. Typically, rely on a GenBank identifier instead of a possibly ambiguous sequence record name. This message is not only to authors, but also to reviewers and editors leveraging resources such as BioSharing^16^, Identifiers.org and MIRIAM [34] repositories in this task. In line with our recommendation, **we propose that publishers provide a dedicated section for obligatory unambiguous references to electronic records**, similar to the traditional bibliographic reference section. This observation echoes recent findings about the lack of clear identification of materials and reagents in scientific papers [35] and recent amendments to data sharing policies by publishers such as PLOS [36].

The second recommendation is **to be explicit about experimental design and experimental variables, identifying the goal of the experiment, independent and response variables**. Table 2 illustrates how variables and sample sizes could be reported in full, allowing a rapid assessment using an *info-box*. Interestingly, as the basic principles of experimental design remain irrespective of the field, it enabled ISA to be applied to non-biological experimental setups as in this case of algorithm comparison. Thinking in terms of experimental design identifies a case of unbalanced factorial design, with study groups of unequal sizes since two bacterial genomes are used but only one genome of mid size and one of large size. Second, it leads one to ask about the state-of-the-art methods for evaluating algorithms to begin with [37, 38] and then for demonstrating process superiority [39]. In the absence of replication for several groups, the estimation of variance and standard deviation cannot be made. Owing to current compute costs, machine availability and project prioritisation, one may consider such a requirement excessive to demonstrate the performance of SOAPdenovo2 when a qualitative assessment may be deemed sufficient. It should, however, be pointed out that from a methodological point of view, applying principles of design of experiments would have certainly emphasised further and demonstrated more compellingly the benefits brought by SOAPdenovo2.

A complementary follow-up to the existing study that would augment it by including additional genomes to collect more data points, thus ensuring replication and balancing of the design. For instance, the *Apis mellifera* (236Mb) genome could be used for the mid-size genome spot and dog (*Canis familiaris*, 2.4 gigabases of haploid genome) genome for the highest size spot. SOAPdenovo2 has been used to assemble the largest animal genome published to date (the 6.5GB *Locust* genome [40]). One could go further still; challenging SOAPdenovo2 and competitors with even larger plant genomes (also notorious for being highly repetitive).

Overall, this second recommendation offers a framework for critical appraisal. The authors conceded that, while the recommendation for testing for more data points along the slope to fully qualify the performance of SOAPdenovo2 algorithm could be justified, the reality of machine occupancy and incurred costs constitute obstacles to effective envelop testing. In addition, *de-novo* sequence assembly of genomes often requires specific parameter tuning to take the specifics of sequence libraries into account (e.g. bacterial artificial chromosome —BAC— or fosmid libraries). Still, those constraints need to be considered and discussed explicitly for the sake of clarity and exhaustivity when reporting results.

A scientific article is a narrative built on results collected through experimentation and facts uncovered through analysis. While the scientific endeavour demands neutrality towards facts, we all know too well the temptation to skew reports to highlight positive results. Hence, the next recommendation is **to remain neutral and report all findings of similar importance with the same weight**. Failing to do so can lead to *jumping to conclusions*, as we witnessed first hand when creating the NPs associated with the SOAPdenovo2 article based on the statements in the abstract.

Three assertions were initially generated: *(A1) increased genome coverage*, *(A2) decreased memory consumption*, *(A3) decreased run time*. Upon verification, *(A3)* turned out to be incorrect. While anecdotal, it is an actual example of *priming* to use Tversky and Kahneman words [41] on the basis of the first two assertions. It also shows a benefit of the NP model, which requires reporting supporting facts back the claims, thus providing a proofing mechanism. Evidence collected from Table 4 in [12] indicated that SOAPdenovo2 took slightly more time when compared to the other two algorithms to reach completion. However, computation run time alone should not be used to assess software performance, as SOAPdenovo2 delivers far more accurate results, as evidenced by the other two variables: *memory consumption* and *genome coverage*. So the metric simply indicates that improvements are often a matter of trade-offs and the observed gains in quality came at the cost of a marginally longer execution. That fact, as measured by the explicit response variable *computation run time* declared in the initial experimental design, should have been stated at the same level as the gain in performance for *genome coverage* and *memory consumption*. The observation confirms the benefits of the declarative aspect of the ISA representation in terms of experimental design, predictor and response variables, as well as the proofing aspect of the NP model.

Thus, the third recommendation can be further specified as **to report all findings corresponding to all the identified response variables**. For its ability to capture provenance and provide attribution, we chose the nanopublication model to report the main findings corresponding to the three response variables in the SOAPdenovo2 experiment, overcoming any priming issue.

Following this model assisted review process which resulted in the identification of a small number of inaccuracies, the authors produced a correction article to officially communicate the amendment to their initial report^17^. Complementing this traditional approach, the release of nanopublications by the present work with the amended values highlights the model’s potential for disseminating evidence.

## Systems and Methods

### 0.7 The ISA model

ISA is a general-purpose metadata tracking framework focused on supporting rich descriptions of the experimental conditions mainly in the Life Sciences, with a growing community of users [6] ranging from institutional [42], project-based and global repositories [43], but also data publication platforms, such as GigaScience and Scientific Data. ISA-Tab is a hierarchical and tabular format designed to represent the experimental design, highlighting both predictor and response variables as well as considering replication of measurements, protocols, procedures and their parameters [44]. At its core, there is an underlying node-edge graph representation where node elements such as *materials* (a cell) and *data* (sequence) are input or outputs of *processes* (*e.g.* purification or data transformation). The ISA-Tab syntax supports use of controlled terminologies or ontologies, also tracking the version and provenance information about those. The ISA open source software suite [26, 44–46] allows for the creation and manipulation of the ISA-Tab formatted information. For this work, the linkedISA software component was used to generate RDF statements from ISA-Tab formatted files, mapping the information to a semantic model and making explicit relations between the entities. In addition, the OntoMaton component [26] was employed to create nanopublications (see section 0.9), which were converted to RDF using NanoMaton^18^.

### 0.8 Galaxy workflow system

The Galaxy project aims to provide software infrastructure enabling scientists to execute complex computational workflows in the field of biology and sequence analysis. It is meant to support data analysis, but also to enable re-enactment and thus reproducibility. Galaxy is an open source, web-based application framework that benefits from a broad user base [8]. In addition to providing executable pipelines in a way that could support the reproduction of the original result, the framework is able to document the process of data analyses by providing a high-level overview diagram of the different analytical steps in the workflow, capturing versions of tools used in analyses and recording intermediate results. An instance of a Galaxy server was set up on GigaScience hardware and Galaxy workflows were defined for each of the algorithms tested.

### 0.9 The Nanopublication model

The Nanopublication (NP) model is a mechanism for enabling the attribution of minimal biological assertions in a machine readable format [47]. Its main components are:

i. the assertion,
ii. the provenance of the assertion,
iii. the publication information of the NP itself, i.e. the attribution of the author(s) of the NP.

The recommended form for exchanging nano publications is by a Semantic Web implementation of the NP minimal model^19^.

### 0.10 The Research Object model

Research Object describes a number of initiatives and approaches trying to describe and associate all of this content together in a machine-readable mechanism so that it can be more easily shared and exchanged. The *researchobject.org* community^20^ involves scientists from a variety of domains to define a principled way for the identification, aggregation and exchange of scholarly information on the Web. It aims to identify the common principles underpinning these various existing solutions in order to create a harmonization of understanding and practices. The Research Object model [10] is one solution among these, providing an aggregation mechanism for components that are constituent parts of a broader research activity. Such components are interrelated with each other and are meant to provide the context to make research more effectively accessible and reusable..

The core RO model is lightweight and domain-neutral, simply providing a bundle structure for aggregating essential information that are needed for reproducing or reusing research results. In this paper, the science workflow-specific Research Object is used, which extends the core Research Object model with workflow-specific terminologies, like the definition of computational workflows, their steps, inputs and outputs data. To create the Research Object presented in this paper, the command-line RO Manager tool [48] was used, which provides the most flexibility for the range of annotations that we could provide. The resulting RO was published in the public RO repository and became accessible at the Research Object Portal [49] through a Permanent Uniform Resource Locator (PURL)^21^.

## Conclusion

The reporting of scientific work can be greatly improved by taking advantage of Galaxy workflows to reenact and validate data analyses in conjunction with the research objects reviewed in this work, namely, ISA, NP and RO data models. They present complementary features, which sweep the entire spectrum of the key points necessary to realise good digital preservation, from ISA and its emphasis on study plans, to the RO model dealing with computational workflow preservation and to NP, harnessed to structure and capture experimental conclusions. The strengths of these complementary data models lie in their respective philosophies. ISA and RO models both provide means to track experimental and computational workflows, with some level of acknowledged overlap which is handled by deferring to the domain specific resources, with the RO project recommending ISA for the biological and life sciences domain.

Yet, it is unrealistic to expect researchers to be deeply acquainted with representation models and other semantic resources. To advance the role for data standards, models and computational workflows in scholarly publishing, we need to make the process viable and above all, scalable. It is therefore critical to re-evaluate the existing tools supporting scholarly publishing. New tools are needed that help navigate and embed semantic representations by integrating representation models seamlessly, vocabulary servers for instance, possibly taking inspiration from NanoMaton^22^, integrating Google’s collaborative spreadsheet environment with ontology lookup and tagging provided by OntoMaton and the NP model. Pivotal to this evolution are the interactions and community liaison needed among a variety of stakeholders, including vocabulary developers, service providers such as BioPortal [27], software developers and publishers, among others. Scholarly publishing has moved to a new phase and will continue to improve as new semantic artefacts are tested in a quest to enhance the article’s content or the discoverability and reuse of the underlying datasets. With peer review costing an estimated 2 billion US dollars each year, and criticisms that it is currently more of a faith rather than evidence-based process [50], this work has at least been an attempt to demonstrate methods of making it a more accurate and quantitative process. Publishers make the argument that they *add value* to the publication process, and these models may offer potential as additional services that can better justify this role.

## Acknowledgments

SAS, PRS and AGB their funding support to BB/L024101/1, FP7 E9RXDC00 (COSMOS), BB/I025840/1 and the University of Oxford e-Research Centre.

The work by PL and SCE and GigaGalaxy and the implementation of workflows were supported from funding from CUHK-BGI Innovation Institute of Trans-omics and School of Biomedical Sciences, The Chinese University of Hong Kong, and China National GeneBank (CNGB).

MSAG and JZ are supported by the EU Wf4Ever project (270129) funded under EU FP7 (ICT-2009.4.1).

MR, MT, EH, RK are supported by the EU Wf4Ever STREP project (270129) funded under EU FP7 (ICT-2009.4.1), the IMI-JU project Open PHACTS (grant agreement n 115191), and RD-Connect (EU FP7/2007-2013, grant agreement No. 305,444).

SOAPdenovo2 was developed with the support of the State Key Development Program for Basic Research of China-973 Program (2011CB809203); National High Technology Research and Development Program of China-863 program (2012AA02A201); the National Natural Science Foundation of China (90612019); the Shenzhen Key Laboratory of Trans-omics Biotechnologies (CXB201108250096A); and the Shenzhen Municipal Government of China (JC201005260191A and CXB201108250096A). Tak-Wah Lam was partially supported by RGC General Research Fund 10612042.

PL, TTL and SCE would like to thank Huayan Gao for technical support on Galaxy. The authors are particularly grateful to Chris Taylor for reading and commenting on the manuscript.

## 1 Authors Contributions

SAS proposed the idea after an initial meeting with JZ, MR, AGB and PRS. SE and PL selected the publication and worked with its authors (RL, TWL). PRS and AGB did ISA-Tab, linkedISA RDF, NPs representation and SPARQL queries over linkedISA and NPs. MR, MT, EH, RK reviewed the NPs. PRS submitted terms to OBI. AGB wrote linkedISA, NanoMaton software and prepared dedicated website and triple store. PL and TLL re-implemented the published SOAPdenovo2 analyses as Galaxy workflows with help from SE, RL and TWL. JZ created the Research Object with input from MSAG and PRS.PRS wrote the manuscript first draft; all authors contributed to the final version, read it and approved it.

## Footnotes

1 GigaScience: http://www.gigasciencejournal.com

2 ScientificData:http://www.nature.com/scientificdata/

3 OECD Principles and Guidelines for Access to Research Data from Public Funding: http://goo.gl/nvi9ME, Berlin Declaration on Open Access to Knowledge in the Sciences and Humanities: http://goo.gl/Kwzo5F, Royal Society-Science as an open enterprise: http://goo.gl/An2X8S

4 GigaDB record: http://dx.doi.org/10.5524/100044

5 SOAPdenovo2 pre-publication history: http://www.gigasciencejournal.com/content/1/1/18/prepub

6 Genome Assembly Gold-standard Evaluations: http://gage.cbcb.umd.edu/

7 GigaGalaxy server: http://galaxy.cbiit.cuhk.edu.hk

8 GigaDB record: http://dx.doi.org/10.5524/100044

9 GAGE analysis script: http://gage.cbcb.umd.edu/results/gage-paper-validation.tar.gz

10 Assembly record: http://www.ncbi.nlm.nih.gov/assembly/2758/

11 PROVenance Ontology (PROV-O): http://www.w3.org/TR/2012/WD-prov-o-20120724/

12 NanoMaton: https://github.com/ISA-tools/NanoMaton

13 STATistics Ontology (STATO): http://stato-ontology.org

14 http://isa-tools.github.io/soapdenovo2/

15 This cost and subsequent calculations were done when the experiment was carried out.

16 http://biosharing.org

17 Correction article: http://goo.gl/LmlpBC

18 NanoMaton: http://github.com/ISA-tools/NanoMaton

19 http://nanopub.org/nschema

20 http://researchobject.org

21 RO PURL: http://goo.gl/14wV3T

22 https://github.com/ISA-tools/NanoMaton

